# Chemical and biophysical characterization of novel potassium channel blocker 3-fluoro-5-methylpyridin-4-amine

**DOI:** 10.1101/2023.08.08.550404

**Authors:** Yang Sun, Sofia Rodríguez-Rangel, Lauren L. Zhang, Jorge E. Sánchez-Rodríguez, Pedro Brugarolas

## Abstract

4-aminopyridine (4AP) is a potassium (K^+^) channel blocker used clinically to improve walking in people with multiple sclerosis (MS). 4AP binds to exposed K^+^ channels in demyelinated axons, reducing the leakage of intracellular K^+^ and enhancing impulse conduction. Multiple derivatives of 4AP capable of blocking K^+^ channels have been reported including three radiolabeled with positron emitting isotopes for imaging demyelinated lesions using positron emission tomography (PET). Here, we describe 3-fluoro-5-methylpyridin-4-amine (5Me3F4AP), a novel K^+^ channel blocker with potential application in PET. 5Me3F4AP has comparable potency to 4AP and the PET tracer 3-fluoro-4-aminopyridine (3F4AP). Compared to 3F4AP, 5Me3F4AP is more lipophilic (logD = 0.664 ± 0.005 *vs.* 0.414 ± 0.002) and slightly more basic (p*K*_a_ = 7.46 ± 0.01 *vs*. 7.37 ± 0.07). In addition, 5Me3F4AP appears to be more permeable to an artificial brain membrane and more stable towards oxidation by the cytochrome P450 enzyme family 2 subfamily E member 1 (CYP2E1), responsible for the metabolism of 4AP and 3F4AP. Taken together, 5Me3F4AP has promising properties for PET imaging warranting additional investigation.

**Significance Statement:** The PET tracer [^18^F]3-fluoro-4-aminopyridine ([^18^F]3F4AP) binds to K^+^ channels in demyelinated axons and has shown promise for imaging demyelinated lesions in animal models. However, its use in humans may be compromised due to rapid metabolism. Thus, a novel 3F4AP derivative amenable to labeling with fluorine-18 was designed and evaluated *in vitro*. The results indicate that 5-methyl-3F4AP exhibits high binding affinity, good physicochemical properties and slower oxidation by CYP2E1 than 3F4AP, making it a promising candidate for further PET studies.

## INTRODUCTION

4-aminopyridine (4AP) is a potassium channel blocker commonly used in the symptomatic treatment of multiple sclerosis (MS) (Stefoski et al., 1987; Goodman et al., 2009; Jensen et al., 2014). Its mechanism of action involves binding from the intracellular side to voltage-gated K^+^ (K_v_) channels exposed due to demyelination, thereby blocking the aberrant efflux of K^+^ ions and enhancing axonal conduction (Sherratt et al., 1980; Bostock et al., 1981; Kirsch et al., 1993; Fehlings and Nashmi, 1996; Rasband et al., 1998; Nashmi and Fehlings, 2001; Devaux et al., 2002; Arroyo et al., 2004; Karimi-Abdolrezaee et al., 2004; Sinha et al., 2006). Additionally, 4AP has demonstrated potential clinical utility for spinal cord injury (Choquet and Korn, 1992; Hayes et al., 1993; Segal et al., 1999; Wolfe et al., 2001; Grijalva et al., 2003; Grijalva et al., 2010), traumatic brain injury (Radomski et al., 2022), and other diseases involving demyelination (Hayes, 2006). Based on the mechanism of action of 4AP, it has been proposed that upregulated K^+^ channels in demyelinated axons could be targeted for imaging demyelination using positron emission tomography (PET) (Brugarolas et al., 2018a; Brugarolas et al., 2018b). Thus, a radiofluorinated derivative of 4AP, [^18^F]3-fluoro-4-aminopyridine ([^18^F]3F4AP), was synthesized and evaluated for imaging demyelination (Brugarolas et al., 2016; Basuli et al., 2018; Brugarolas et al., 2018b). In those studies, [^18^F]3F4AP displayed high sensitivity in detecting demyelinated lesions in rodent models of MS (Brugarolas et al., 2018b) and non-human primates (Guehl et al., 2021b). Furthermore, [^18^F]3F4AP has shown acceptable radiation dosimetry in healthy human volunteers (Brugarolas et al., 2022) and it is currently undergoing evaluation in MS patients (Guehl et al., 2022) (ClinicalTrials.gov identifier: NCT04699747). Nevertheless, in humans [^18^F]3F4AP has shown lower metabolic stability (<50% parent fraction (PPF) remaining, 30 min post injection (Brugarolas et al., 2022)) than in monkeys (>90% PPF 2 h post injection (Guehl et al., 2021b)), which could potentially make its quantification challenging. This reduction in metabolic stability was found to arise from the inhibitory effect of isoflurane on the metabolism of [^18^F]3F4AP as evidenced by the fact that awake mice metabolize the tracer faster than anesthetized mice, with 20 ± 4 PPF *vs*. 65 ± 7 PPF respectively 35 min post-injection (Ramos-Torres et al., 2022). Additional studies indicate that [^18^F]3F4AP is oxidized at the 5-position to 5-hydroxy-3F4AP by the cytochrome P450 enzyme CYP2E1 (Sun et al., 2023), which prompted us to look for more stable derivatives with suitable binding affinity and brain permeability.

Studies on 4AP derivatives including 3,4-diaminopyridine, 4-aminopyridine-3-methanol and others demonstrate that small substituents in the 3-position do not significantly impair binding to K^+^ channels (Kirsch and Narahashi, 1978; Berger et al., 1989; Caballero et al., 2007; Sun et al., 2009; Brugarolas et al., 2018b; Rodriguez-Rangel et al., 2020). Based on this, several 4AP derivatives labeled with carbon-11, namely [^11^C]3-trifluoromethyl-4AP (Ramos-Torres et al., 2020), [^11^C]3-methoxy-4AP (Guehl et al., 2021a), and [^11^C]3-methyl-4AP (Sun et al., 2022), have also been investigated (**Table 1**). Although some of these derivatives possess advantages such as higher binding affinity and specific binding compared to [^18^F]3F4AP (Ramos-Torres et al., 2020), fluorine-18 labeled tracers are generally preferred due to their longer half-life (110 min *vs.* 20.3 min) (Pike, 2009). Furthermore, [^18^F]3F4AP displays excellent characteristics for PET imaging including a fast entry and washout from the brain mediated by a p*K*_a_ value close to physiological (p*K*_a_ = 7.65) and a positive logD (logD = 0.41) (Rodriguez-Rangel et al., 2020). In addition, recent studies have shown that 3-methyl-4AP has good binding affinity towards K^+^ channels, and thus, we hypothesized that a derivative of 3F4AP with a methyl group at the 5-position, 5Me3F4AP, would retain its ability to block K_v_ channels and have adequate brain permeability while exhibiting improved stability against metabolism. Herein, we characterized the pharmacological and biophysical properties of 5Me3F4AP and evaluated its *in vitro* metabolic stability towards CYP2E1 to investigate its potential as a PET tracer for imaging demyelination.

**Table 1.**
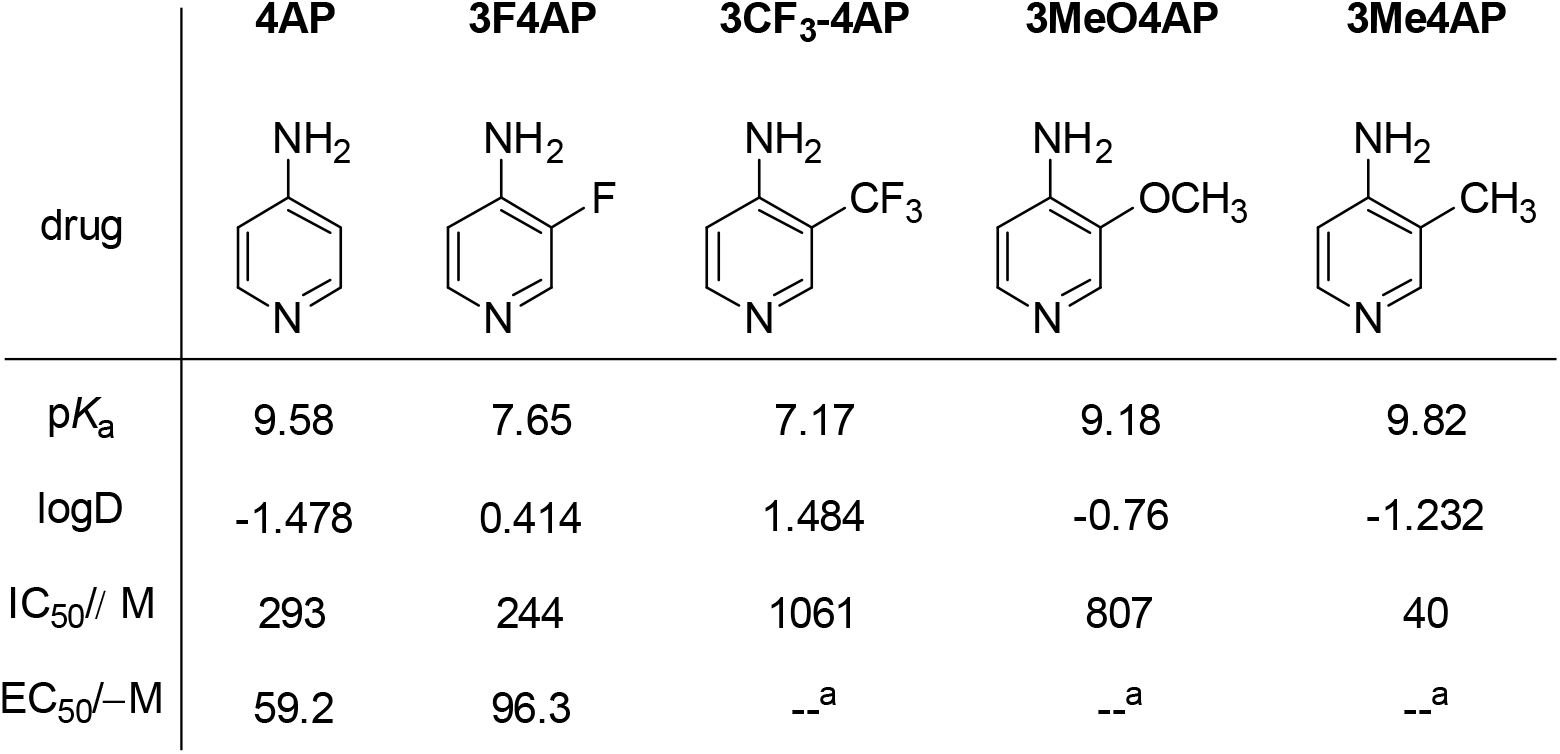
Structures of 4AP, 3F4AP, 3CF_3_4AP, 3MeO4AP, 3Me4AP and their p*K*_a_, logD, K_v_1 IC_50_ and EC_50_ values.

|  | 4AP | 3F4AP | 3CF<sub>3</sub>-4AP | 3MeO4AP | 3Me4AP |
| --- | --- | --- | --- | --- | --- |
| drug |  |  |  |  |  |
| p*K*<sub>a</sub> | 9.58 | 7.65 | 7.17 | 9.18 | 9.82 |
| log*D* | -1.478 | 0.414 | 1.484 | -0.76 | -1.232 |
| IC<sub>50</sub>/M | 293 | 244 | 1061 | 807 | 40 |
| EC<sub>50</sub>/M | 59.2 | 96.3 | --<sup>a</sup> | --<sup>a</sup> | --<sup>a</sup> |

## MATERIALS AND METHODS

### Animal Studies Compliance

Methods involving *Xenopus laevis* frogs were performed in accordance with relevant guidelines and regulations and with the approval of the Comité Institucional del Cuidado y Uso de Animales en el Laboratorio at the University of Guadalajara (protocol: CUCEI/CINV/CICUAL-03/2023).

### Materials

5Me3F4AP was purchased from AmBeed Inc. (Arlington Hts, IL, USA). All other chemical compounds used for this study were purchased from Sigma-Aldrich Merck (Merck KGaA, Darmstadt, Germany) or as otherwise indicated.

### Partition coefficient determination

The octanol-water partition coefficient (logD) at pH 7.4 was determined according to our previous reported protocol (Rodriguez-Rangel et al., 2020). Briefly, PBS (900 μL), 1-octanol (900 μL), and a 10 mg/ mL aqueous solution of each compound (2 μL) were added to a 2 mL HPLC vial. The compounds were partitioned between the layers via vortexing and centrifuged at 1,000 g for 1 min to allow for phase separation. A 10 μL portion was taken from each layer (autoinjector was set up to draw volume at two different heights) and analyzed by HPLC. The relative concentration in each phase was determined by integrating the area under each peak and comparing the ratio of the areas from the octanol and aqueous layers. A calibration curve was performed to ensure that the concentrations detected were within the linear range of the detector (see **Figure S1 and S2** in Supporting Information). This procedure was repeated four times for each compound.

### Determination of pK_a_

The p*K*_a_ was determined using titration according to our previously described protocol (Rodriguez-Rangel et al., 2020). A 1 mg/mL solution of 5Me3F4AP was prepared, of which 5 ml was titrated with 0.01M HCl solution beyond the equivalence point. After each incremental addition of titrant, the sample was stirred and the pH reading was taken with a pH meter. The Gran plot of the titration was analyzed to obtain the p*K*_a_ (see the plot in Supporting Information, **Figure S3**). A similar protocol was used to titrate 3F4AP and 4AP, respectively (**Figures S4** and **S5**). The titration was repeated four times each for each compound.

### Permeability rate determination

The permeability rates of 5Me3F4AP and 3F4AP were determined using Parallel Artificial Membrane Permeability Assay-blood-brain barrier (BBB) kit (BioAssay Systems, Hayward, USA) following the manufacturer’s protocol. Initially, solutions of each test compound were prepared in DMSO at a concentration of 10 mM. These stock solutions along with the stock solutions of control compounds (high control: promazine hydrochloride, low control: diclofenac) were then diluted with PBS (pH = 7.2) to obtain the donor solutions with a final concentration of 500 μM. At the same time, 200 μM of equilibrium standards for each compound and a DMSO blank control solution were prepared.

In the experimental setup, 300 μL of PBS was added to the desired well of the acceptor plate, and 5 μL of BBB lipid solution in dodecane was added to membranes of the donor plate. Next, 200 μL of the donor solutions of each test compound and each permeability control were added to the duplicate wells of the donor plate. The donor plate was carefully placed on the acceptor plate and incubator for 18 hours at room temperature. After incubation, UV absorption measurements were conducted using 100 μL of the resulting solutions from the acceptor plate and the equilibrium standards. UV absorption of the controls was measured by running a UV scan in the range of 200 to 500 nm. UV absorption of 5Me3F4AP and 3F4AP was measured using HPLC equipped with a UV detector and C18 column. The calibration curve, demonstrating the relationship between the area under the curve (AUC) in the HPLC chromatogram and concentration, is presented in the Supporting Information (**Figure S1, S2**).

### Cut-Open Voltage Clamp Electrophysiology

Blocking potency of 5Me3F4AP was evaluated on the voltage-gated Shaker (homologous to mammalian K_v_1.2) ion channel expressed in *Xenopus laevis* oocytes, as previously described (Rodriguez-Rangel et al., 2020). Briefly, each oocyte expressing the Shaker channel was voltage-clamped in a Cut-Open Voltage Clamp (COVC) station in order to elicit K^+^ currents in response to the voltage stimulus protocol, which entailed steps of 50 ms from -100 to 60 mV in increments of 10 mV. External and internal recording solutions for COVC were composed (in mM) of 12 KOH, 2 Ca(OH)_2_, 105 NMDG-MES, 20 HEPES and 120 KOH, 2 EGTA, and 20 HEPES, respectively, with pH adjusted to 7.4 with methylsufonate. For measurements achieved at pH = 9.1 or 6.4, HEPES was replaced by 2-(cyclohexylamino)-ethanesulfonic acid or 2-(N-Morpholino)ethanesulfonic acid, respectively. Each oocyte expressing the Shaker ion channel was voltage-clamped to record K^+^ currents, first in the absence of 5Me3F4AP, and subsequently with the addition of 5Me3F4AP, from 0.0001 to 10 mM. Relative current (I_rel_) was quantified as the ratio of the current in the absence and in the presence of the indicated concentration of 5Me3F4AP. Finally, K^+^ currents were amplified with the Oocyte Clamp Amplifier CA-1A (Dagan Corporation, Minneapolis, MN, USA) and digitized with the USB-1604-HS-2AO Multifunction Card (Measurement Computing, Norton, MA, USA). All systems were controlled with the GpatchMC64 program (Department of Anesthesiology, UCLA, Los Angeles, CA, USA) via a PC. Electrophysiology recordings were sampled at 100 kHz and filtered at 10 kHz.

### Electrophysiology data analysis

Data analysis was performed as previously described (Rodriguez-Rangel et al., 2020). Briefly, the half-maximal inhibitory concentration of 5Me3F4AP (IC_50_) was determined by fitting the I_rel_ curve to the Hill equation at each value of V and pH. A Hill coefficient (h) in the range of 0.9<h<1.1 was used. Voltage and pH dependence of IC_50_ was analyzed by fitting the IC_50_(V) at each pH with a one-step model of inhibition (Woodhull model) which allowed the determination of the fractional distance through the membrane electrical field (δ) that 5Me3F4AP has to cross to reach its binding site (Woodhull, 1973):

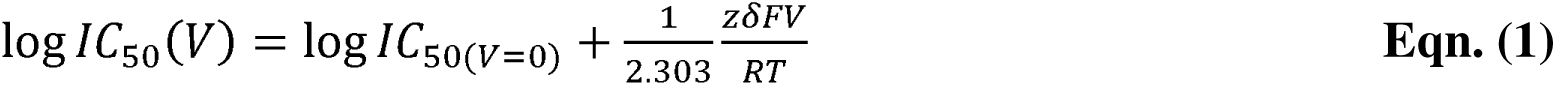

where IC_50(V= 0)_ is the value of IC_50_ at V = 0 mV, F is the Faraday constant, R is the gas constant, T is the room temperature, and z is the apparent charge.

Mean values of data ± standard deviation (s.d.) are given or plotted and the number of experiments is denoted by n. Upper and lower limits of the 95% of confidence interval (CI_95_) are denoted as 10^(log *IC*50+*s.d*)^ and10^(log *IC*50-*s.d*)^, respectively.

### CYP2E1-mediated metabolic stability assessment

The relative metabolic stability towards CYP2E1 was assessed with the competitive CYP2E1 inhibition assay utilizing the Life Technologies^TM^ Vivid^®^ CYP2E1 screening kit, as described in previous studies (Sun et al., 2023). In this assay, fluorescence emitted by the metabolic product of a specific CYP2E1 substrate included in the kit, was measured in the absence and presence of substrate competitors. Consequently, the highest fluorescence values were obtained from the blank experiments, lacking any competitors. As the concentration of competitors increased, or more potent competitors were introduced, the fluorogenic emission decreased accordingly. Specifically, 40 µL of 2.5X (final concentration 25µM) solution of test compounds (4AP, 3F4AP, 5Me3F4AP, and positive control, i.e., tranylcypromine) in 1X Vivid^®^ CYP2E1 reaction buffer was added to desired wells of a falcon black/clear 384-well plate in three replicates. Afterwards, 50 µL master pre-mix 2X (40 nM) CYP2E1 BACULOSOMES^®^ and 2X (0.6 Units/mL) Vivid^®^ regeneration system in 1X reaction buffer) was added to each well. The plate was incubated for 10 minutes at room temperature to allow the compounds to interact with the CYP2E1 in the absence of enzyme turnover. Next, the reaction was initiated by adding 10µL per well of 10X (100 μM) Vivid^®^ substrate (2H-1-benzopyran-3-carbonitrile,7-(ethoxy-methoxy)-2-oxo-(9Cl)) and 10X (300 μM) Vivid® NADP^+^ mixture. Immediately (in less than 2 minutes), the plate was transferred into the fluorescent plate reader and fluorescence was monitored over 60 minutes (reads in 1-minute intervals) at 415 nm as excitation wavelength and 460 nm as emission wavelength. The obtained reads were plotted using GraphPad Prism 9.

### Determination of the IC_50_ of 5Me3F4AP to CYP2E1

A similar Vivid^®^ CYP2E1 assay was conducted as described above. Instead of testing a single concentration of 5Me3F4AP (final concentration 15µM), a series of concentrations (4.0 mM, 1.2 mM, 400 µM, 120 µM, 40 µM, 12 µM, 4.0 µM, 1.2 µM) were tested with three replicates for each concentration. The plate fluorescence was monitored over 60 minutes (reads in 1-minute intervals) at 415 nm as excitation wavelength and 460 nm as emission wavelength. The reads at 60 min (recalculated by the linear trend line equation) of each concentration were used and fitted with GraphPad Prism9 dose-response-inhibition (concentration is log) curve fitting to calculate the IC_50_ values. A similar procedure was used for determining the IC_50_ of 3F4AP.

## Data availability

The datasets generated during and/or analyzed during the current study are available from the corresponding authors upon request.

## RESULTS

### Basicity, lipophilicity and membrane permeability of 5Me3F4AP

Measurements of these pharmacological parameters of 5Me3F4AP were taken and compared to those of its predecessors (**Table 2**). As indicated in column 2, 5Me3F4AP demonstrates slightly higher basicity in comparison to 3F4AP (7.46 ± 0.01 *vs.* 7.37 ± 0.07). Both compounds have p*K*_a_ values that are close to the physiological pH, indicating their coexistence in both protonated and neutral forms under physiological conditions. In contrast, 4AP and 3Me4AP display greater basicity (p*K*_a_ values above 9), indicating that the protonated form is predominant at physiological pH.

**Table 2.** Pharmacological parameters for 5Me3F4AP, 3F4AP, 4AP and 3Me4AP.

| Drug | $pK_a$ | logD<br>(pH 7.4) | $P_e$<br>(nm/s) |
| --- | --- | --- | --- |
| 5Me3F4AP | $7.46 \pm 0.01$ | $0.664 \pm 0.005$ | $43.3 \pm 0.5$ |
| 4AP | $9.19 \pm 0.03$ | $-1.478 \pm 0.014^a$ | $2.36 \pm 0.03^b$ |
| 3F4AP | $7.37 \pm 0.07$ | $0.414 \pm 0.002^a$ | $13.6 \pm 0.5^c$ |
| 3Me4AP | $9.82 \pm 0.06^a$ | $-1.232 \pm 0.008^a$ | -- |

logD: experimental partition coefficient octanol: water at pH 7.4 (n =4).
$pK_a$ : experimental titration coefficient at room temperature (n =3).
$P_e$ : permeability rate across the artificial membrane (n = 2).
<sup>a</sup> previously reported data using the equipment and protocol (Rodriguez-Rangel et al., 2020).
<sup>b</sup> previously reported data (Brugarolas et al., 2018b).
<sup>c</sup> result obtained during the present study, a value of $15.6 \pm 0.6$ nm/s was previously reported (Brugarolas et al., 2018b).

In terms of lipophilicity, 5Me3F4AP shows an octanol/water partition coefficient value at pH 7.4 of 0.664 ± 0.005 (**Table 2. Column 3**), which is higher than that of 3F4AP (logD = 0.414 ± 0.002). This result indicates that both compounds preferentially partition into the octanol layer, potentially facilitating faster permeation through a lipophilic membrane like the BBB via passive diffusion. Conversely, 4AP and 3Me4AP exhibit a preference for partitioning in the water layer (logD_4AP_ = -1.478 ± 0.014, logD_3Me4AP_ = -1.232 ± 0.008), suggesting slower permeation rates. This trend was further validated through a parallel artificial membrane permeability assay, which demonstrated that 5Me3F4AP permeates approximately three times faster than 3F4AP and 18 times faster than 4AP (**Table 2. Column 4**).

### Affinity towards K^+^ channels

The blocking potency at different pH conditions (6.4, 7.4 and 9.1) and voltages (-100 to 60 mV) of 5Me3F4AP was evaluated by measuring the K^+^ currents generated by Shaker voltage-gated potassium channel from *D. melanogaster* heterologously expressed in *Xenopus laevis* oocytes (**Figure 1**). Specifically, **Figure 1A** shows three representative recordings elicited as a response to the voltage stimulus before and after the application of 1 mM of 5Me3F4AP under three extracellular pH conditions. From these recordings, it is clear that 1 mM 5Me3F4AP efficiently blocks the channels. **Figure 1B** shows the dose-response plot (relative K^+^ currents at 40 mV *vs*. 5Me3F4AP concentration) and the fitting of the experimental values to Hill equation to calculate IC_50_. This plot shows that blocking is dose and pH dependent with less efficient blocking at higher pH. **Figure 2C** shows the calculated IC_50_ values at different values of pH. A model of electric field constant (Woodhull model, **Equation 1**) was used to determine their biophysical parameters IC_50(at V=0mV)_ and the electric fraction (δ), which represents the fraction of the electric field that the compound must cross through the channel pore to reach its binding site. This plot shows that the IC_50_ of 5Me3F4AP increases with voltage and pH indicating a drop in potency. IC_50_ and δ values determined from fitting the Hill and Woodhull equations are shown in **Table 3** and compared to those of 4AP, 3F4AP, and 3Me4AP. These values confirm that at pH 7.4, the blocking potency of 5Me3F4AP to K_v_ channels is similar to that of 4AP and 3F4AP previously reported (Brugarolas et al., 2018b; Rodriguez-Rangel et al., 2020). In addition, when the pH was increased from 6.4 to 9.1, the IC_50_ of 5Me3F4AP (and 3F4AP) increased around 3-fold indicating lower potency. Conversely, the IC_50_ from 4AP and 3Me4AP decreased by approximately 4- and 3-fold, respectively, indicating higher potency at higher pH. These differences in pH dependence confirm that it is the protonated form that preferentially binds to channel, since 4AP and 3Me4AP exist mostly in their protonated state at this pH range, whereas 3F4AP and 5Me3F4AP are mostly protonated at pH 6.4 and predominantly neutral at pH 9.1. Finally, a δ value of ∼0.4 was obtained for 5Me3F4AP; this value is consistent with the δ values of 4AP, 3F4AP and 3Me4AP previously reported (Rodriguez-Rangel et al., 2020), and it indicates that this novel blocker binds at the same site within the Shaker pore traversing ∼ 40% of the electric field generated across the lipid bilayer. Therefore, these findings demonstrate that the blocking potencies of these K_V_ Shaker-related channel blockers produce a trend as follows: 3Me4AP > 3F4AP ∼ 4AP ∼ Me3F4AP.

**Figure 1.**
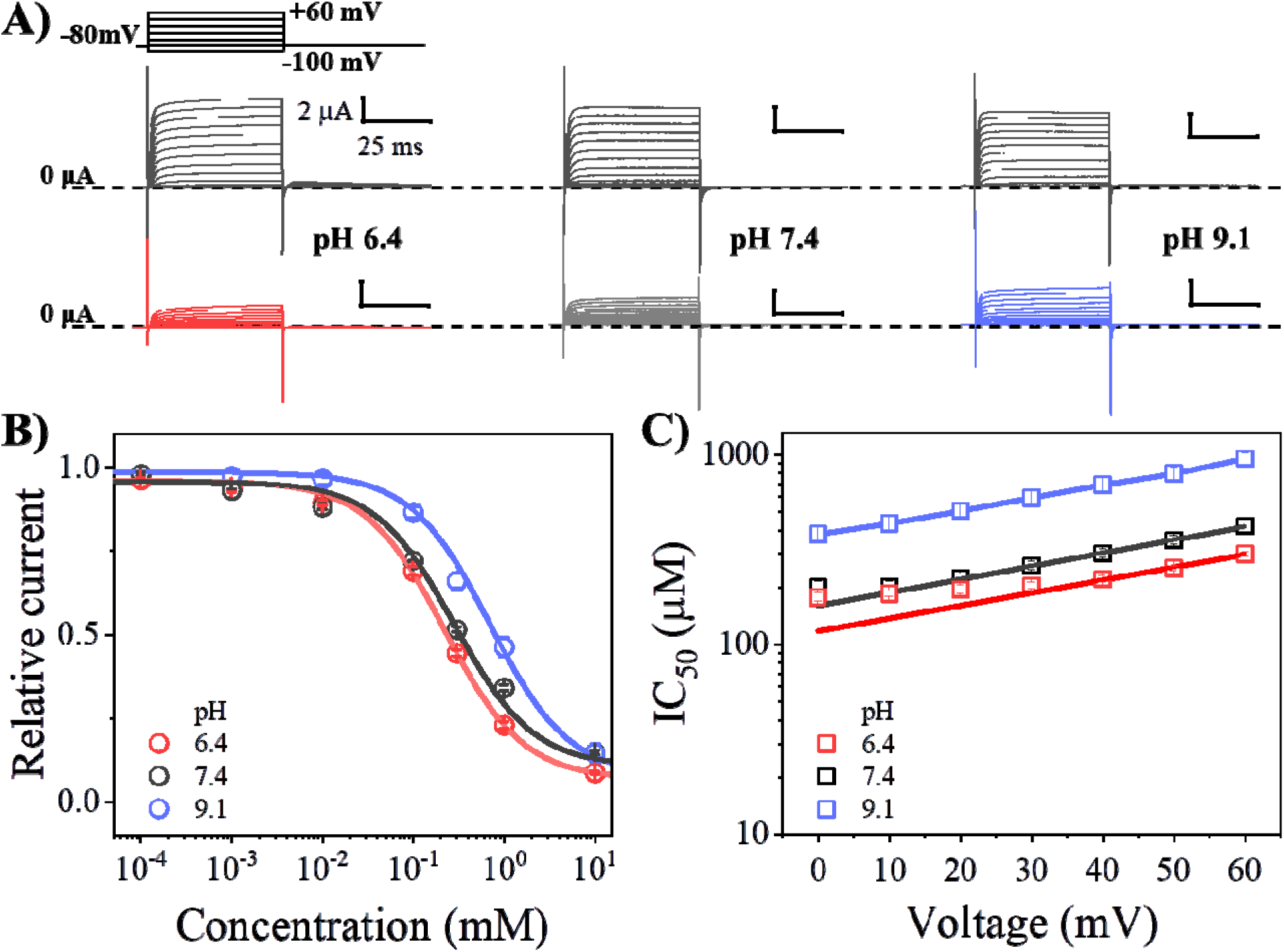
Pharmacological and biophysical characterization of 5Me3F4AP upon Shaker K_V_ ion channel. (**A**), Representative recordings at pH values of 6.8, 7.4 and 9.1 elicited from three different oocytes expressing the Shaker channel before (upper, black) and after (lower, colored) the blockage with 1 mM of 5Me3F4AP. Currents were recorded as the response to voltage stimulus protocol that consisted of 50 ms depolarization steps from -100 to 60 mV in increments of 10 mV (top left). Dashed line represents the zero current value. Horizontal and vertical bars of 25 ms and 2 μA represent the time and current scale for all recordings. (**B**), Relative current *vs.* concentration of 5Me3F4AP curves assessed at 40 mV and (**C**), IC_50_ *vs.* voltage curves at different pH values. Continuous lines of panels **B** and **C** represent the fits with the Hill equation and Woodhull model (**Equation 1**), respectively. Fit parameters are given in **Table 3**.

**Figure 2.**
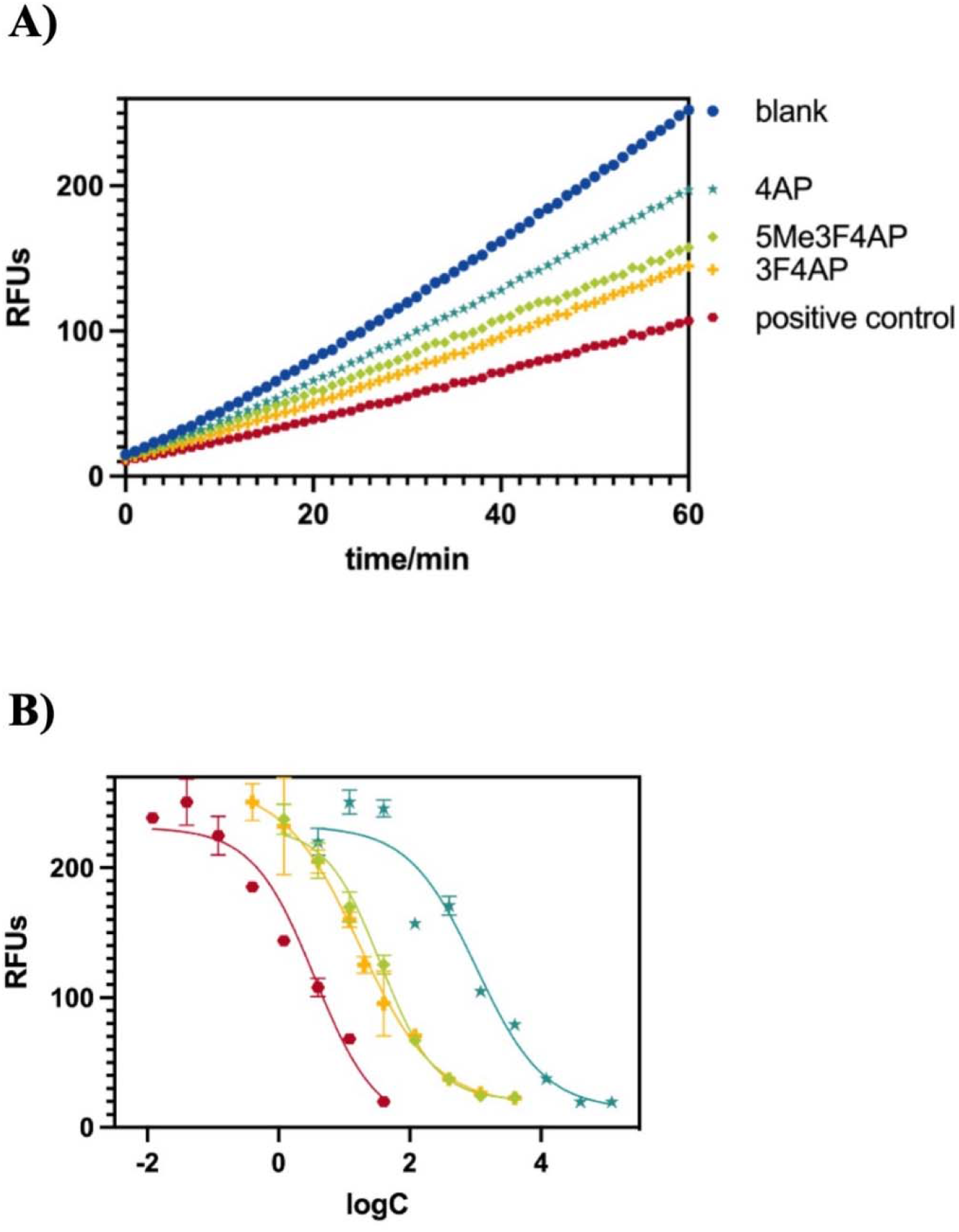
Competitive inhibition of CYP2E1. (Left) Relative fluorescence-time curves based on the 60-minute kinetic measurement of 4AP (cyan), 5Me3F4AP (green), 3F4AP (yellow), and tranylcypromine (red, positive control). (Right) IC_50_ fit curves for 4AP, 5Me3F4AP, 3F4AP, and tranylcypromine with the same color sets.

**Table 3.**
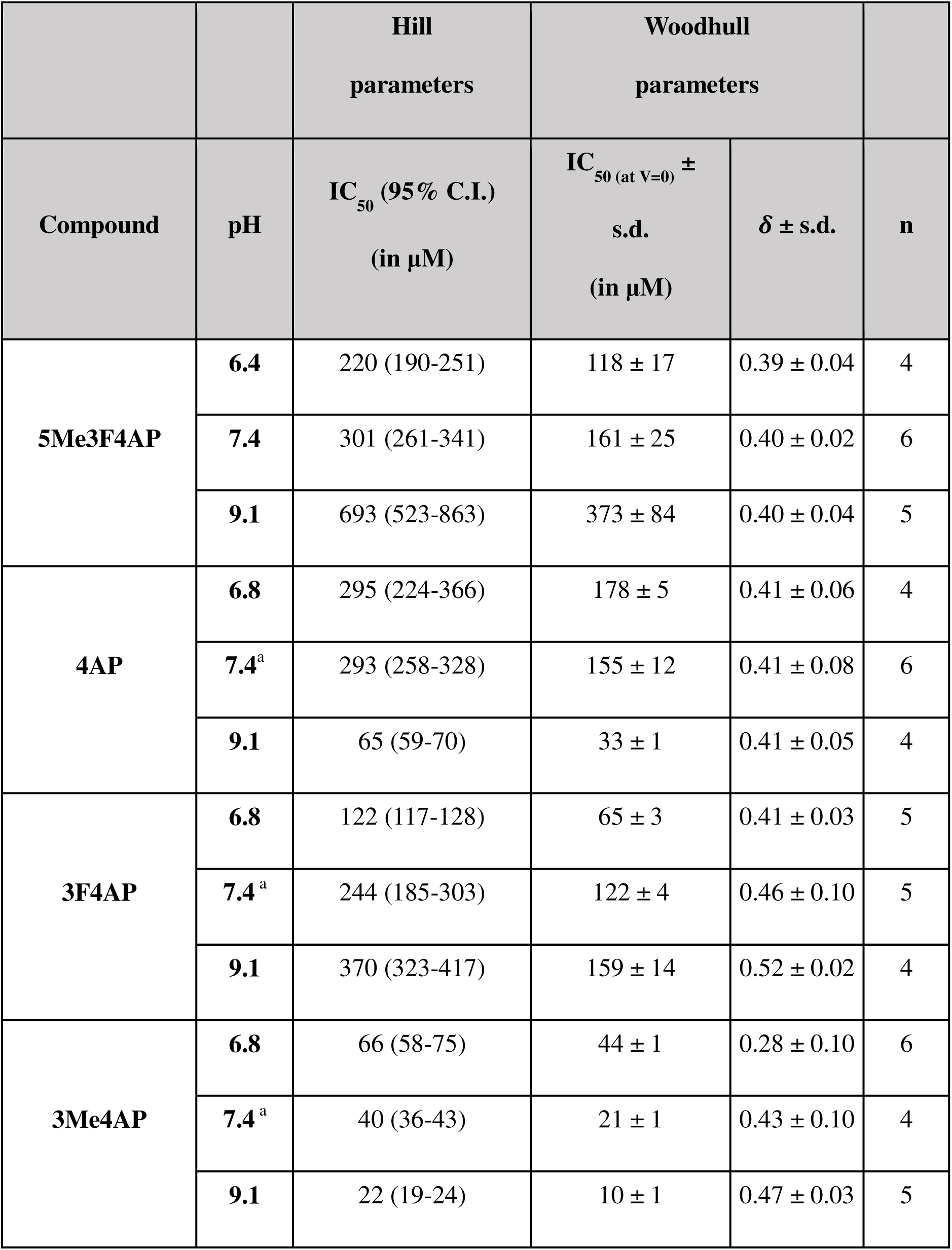
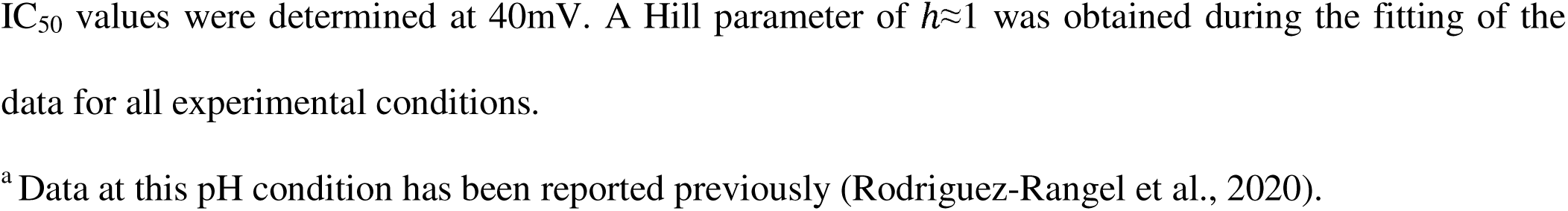
IC_50_ values of 5Me3F4AP: Hill and Woodhull parameters.

|  |  | Hill<br>parameters | Woodhull<br>parameters |  |  |
| --- | --- | --- | --- | --- | --- |
| Compound | pH | IC<sub>50</sub> (95% C.I.)<br>(in μM) | IC<sub>50</sub> (at V=0) ±<br>s.d.<br>(in μM) | $\delta \pm$ s.d. | n |
| 5Me3F4AP | 6.4 | 220 (190-251) | 118 ± 17 | 0.39 ± 0.04 | 4 |
| | 7.4 | 301 (261-341) | 161 ± 25 | 0.40 ± 0.02 | 6 |
| | 9.1 | 693 (523-863) | 373 ± 84 | 0.40 ± 0.04 | 5 |
| 4AP | 6.8 | 295 (224-366) | 178 ± 5 | 0.41 ± 0.06 | 4 |
| | 7.4<sup>a</sup> | 293 (258-328) | 155 ± 12 | 0.41 ± 0.08 | 6 |
| | 9.1 | 65 (59-70) | 33 ± 1 | 0.41 ± 0.05 | 4 |
| 3F4AP | 6.8 | 122 (117-128) | 65 ± 3 | 0.41 ± 0.03 | 5 |
| | 7.4<sup>a</sup> | 244 (185-303) | 122 ± 4 | 0.46 ± 0.10 | 5 |
| | 9.1 | 370 (323-417) | 159 ± 14 | 0.52 ± 0.02 | 4 |
| 3Me4AP | 6.8 | 66 (58-75) | 44 ± 1 | 0.28 ± 0.10 | 6 |
| | 7.4<sup>a</sup> | 40 (36-43) | 21 ± 1 | 0.43 ± 0.10 | 4 |
| | 9.1 | 22 (19-24) | 10 ± 1 | 0.47 ± 0.03 | 5 |

IC<sub>50</sub> values were determined at 40mV. A Hill parameter of $h \approx 1$ was obtained during the fitting of the data for all experimental conditions.
<sup>a</sup> Data at this pH condition has been reported previously (Rodriguez-Rangel et al., 2020).

### Metabolic stability towards CYP2E1 of 5Me3F4AP

To estimate the metabolic stability of 5Me3F4AP towards CYP2E1, we conducted an *in vitro* investigation utilizing a competitive inhibition assay. The protocol used for this study followed a previously established method (Marks et al., 2002). According to the principle of this assay outlined in the method section, compounds that are good substrates of CYP2E1 result in greater reduction in the rate of formation of a fluorescent reporter than compounds that are poor substrates. In this study, we measured the reaction rates without competitor (blank) as well as in the presence of tranylcypromine (positive control), 4AP, 3F4AP and 5Me3F4AP. As illustrated in **Figure 2**, the addition of tranylcypromine, a widely recognized potent substrate of CYP2E1, resulted in the most pronounced reduction in fluorogenic emission when compared to the reaction conducted without any addition of enzyme substrates (red *vs*. blue lines). In comparison, 4AP exhibited a minor reduction in fluorogenic emission (cyan *vs.* blue lines) indicating that it is a poor substrate of CYP2E1. 3F4AP demonstrated a substantial decrease in rate (yellow *vs.* blue lines) indicating 3F4AP is a good substrate of CYP2E1, undergoing metabolism at a much faster rate than 4AP. In comparison, 5Me3F4AP demonstrated a reaction rate between 4AP and 3F4AP, bearing a higher resemblance to 3F4AP (**Figure 2**. green *vs.* cyan and yellow lines). To further quantify the inhibition potency of 5Me3F4AP towards CYP2E1, we measured the CYP2E1-mediated reaction rate in the presence of varying concentrations of 5Me3F4AP and 3F4AP and performed the dose-response fitting. We also compared these results with our previous results for 4AP, 3F4AP and the positive control tranylcypromine (Sun et al., 2023). This analysis showed that 5Me3F4AP has an IC_50_ about two times higher than 3F4AP and about 23 times lower than 4AP (**Table 4**, entries 1 to 3), indicating that it is more stable than 3F4AP but not as stable as 4AP.

**Table 4.** IC_50_ values and their respective confidence interval (C.I.) for CYP2E1 substrates.

| Entry | Drug | IC<sub>50</sub> (μM) | 95% C.I. (μM) | n |
| --- | --- | --- | --- | --- |
| 1 | 5Me3F4AP | 35.9 | 26.2-45.9 | 3 |
| 2 | 4AP<sup>a</sup> | 831 | 605-1199 | 3 |
| 3 | 3F4AP | 17.0 | 13.9-25.2 | 6 |
| 4 | Tranylcypromine<sup>a</sup> | 2.3 | 1.5-3.4 | 3 |

<sup>a</sup> Previously reported data (Sun et al., 2023).

## DISCUSSION

This study identified 5Me3F4AP as novel K^+^ channel blocker with potential application for PET and examined critical pharmacological properties including lipophilicity, basicity, membrane permeability, target binding affinity and metabolic stability. In comparison to its predecessor 3F4AP, 5Me3F4AP was found to have higher lipophilicity (logD of 0.66 *vs.* 0.41) and slightly higher basicity (p*K*_a_ 7.46 *vs.* 7.37). The lower p*K*_a_ values of 3F4AP and 5Me3F4AP compared to 4AP (p*K*_a_ 9.58) indicate that these compounds exist both in the neutral and protonated form at physiological pH, which facilitates the entry into the brain since only the neutral form can cross the BBB through passive diffusion. This effect was confirmed using a membrane permeability assay which showed that 5Me3F4AP cross an artificial brain membrane approximately 3 times faster than 3F4AP. These findings suggest that 5Me3F4AP will have higher brain uptake than 3F4AP.

Regarding blocking potency, it was found that at pH 7.4 the potency of 5Me3F4AP is very similar to that of 4AP and 3F4AP. At higher pH the potency of 3F4AP and 5Me3F4AP dropped significantly but not that of 4AP supporting that only the protonated form is able to block the channel (Choquet and Korn, 1992) since 3F4AP and 5Me3F4AP exist predominantly in the neutral form at basic pH. These findings suggest that 5Me3F4AP will bind to demyelinated lesions with similar sensitivity as 3F4AP.

In terms of metabolic stability, 5Me3F4AP was found to undergo CYP2E1-mediated oxidation about two times slower than 3F4AP (**Figure 2**, green *vs.* yellow lines), which was further confirmed by calculating the IC_50_ (35.9 *vs.* 17.0). Although the oxidation rate of 5Me3F4AP was still significantly faster than 4AP (**Figure 2**, green *vs.* cyan lines), this reduction in the rate of oxidation suggest that 5Me3F4AP will have greater *in vivo* stability than 3F4AP. A limitation of this study is that we did not test other possible metabolic enzymes, which may play a role *in vivo*. Nevertheless, it is reasonable to test CYP2E1 given that prior studies strongly suggest this enzyme is primarily responsible for the metabolism of 3F4AP and 4AP (Caggiano and Blight, 2013; Brugarolas et al., 2022)) and the *in vivo* stability of 4AP (∼70% parent fraction in plasma 24 h post oral administration (Caggiano and Blight, 2013) compared to 3F4AP (< 50% parent fraction in plasma 60 min post intravenous administration (Brugarolas et al., 2022)) is consistent with the measured CYP2E1 oxidation rates for these compounds. In sum, the favorable characteristics of 5Me3F4AP position it as an intriguing candidate worthy of further investigation as a potential alternative to [^18^F]3F4AP for PET imaging.

## Abbreviations

4AP: 4-aminopyridine;
AUC: area under curve;
BBB: blood-brain barrier;
CI_95_: 95% of confidence interval
COVC: Cut-Open Voltage Clamp;
CYP2E1: cytochrome P450 family 2 subfamily E member 1;
EGTA: ethylene glycol-bis(**ß**-aminoethyl ether)-*N,N,N’,N’*-tetraacetic acid;
3F4AP: 3-fluoro-4-aminopyridine;
HEPES: *N*-2-hydroxyethyl-piperazine-*N’*-2-ethanesulfonic acid;
HPLC: high-performance liquid chromatography;
HRMS: high resolution mass spectrometry;
IC_50_: half maximal inhibitory concentration;
5Me3F4AP: 3-fluoro-5-methylpyridin-4-amine;
MES: 2(*N*-morpholino)ethanesulfonic acid;
MS: multiple sclerosis;
NMDG: *N*-methyl-D-glucamine;
3OH4AP: 3-hydroxy-4-aminopyridine;
PBS: phosphate buffered saline;
PET: positron emission tomography;
PPF: percent of parent fraction;
RFUs: relative fluorescence units;
s.d.: standard deviation.

## Section

Drug Discovery and Translational Medicine

## Author contributions

Y.S. performed the logD, artificial membrane permeability and CYP2E1 metabolic stability experiments and analyzed the data. S.R.R performed the COVC experiments and analyzed the data. L.L.Z. performed the p*K*_a_ experiments and analyzed the data. P.B. and J.E.S.R. conceived and supervised the project. All authors wrote the manuscript.

## Footnotes

This study was supported by the National Institute of Neurological Disorders and Stroke Grant R01NS114066 (P.B.); PROSNI-UdeG 2022, Mexico (J.E.S.R.) and CONAHCyT, Mexico (886951) (S.R.R.).

