## Supplemental information for "Chemical and biophysical characterization of novel potassium channel blocker 3-fluoro-5-methylpyridin-4-amine"

for

**Figure S3.** Gran plot of 5Me3F4AP in titration with hydrochloric acid (volume in mL)……………………………S4

**Figure S4.** Gran plot of 3F4AP in titration with hydrochloric acid (volume in mL)……………..……………………S4

**Figure S5.** Gran plot of 4AP in titration with hydrochloric acid (volume in mL)………………………………………S5

| logC | AUC |
| --- | --- |
| 4 | 2.84832 |
| 3.47712 | 2.33759 |
| 3 | 1.86438 |
| 2.47712 | 1.3472 |
| 2 | 0.870304 |
| 1.47712 | 0.346001 |

### **Figure S1.** Calibration curves of 5Me3F4AP in analytical HPLC at 254 nm (concentration in μM).

| logm | AUC |
| --- | --- |
| 1 | 2.60519 |
| 0.477121 | 2.08992 |
| 0 | 1.61237 |
| -0.522879 | 1.08622 |
| -1 | 0.603891 |
| -1.52288 | 0.0576281 |
| -2 | -0.413976 |

### **Figure S2.** Calibration curves of 3F4AP in analytical HPLC at 254 nm (mass in μg).

**Figure S3.** Gran plot of 5Me3F4AP titrated with HCl.

5Me3F4AP Gran plot

Vi * 10pH

| Vi (ml) | Run 1 | Run 2 | Run 3 | Run 4 |
| --- | --- | --- | --- | --- |
| 1 | 80233222 | 72926838 | 77946412 | 73732149 |
| 2 | 41195299 | 42064579 | 44219501 | 36546953 |
| 2.5 | 28664710 | 27505840 | 26565424 | 21861180 |
| 3 | 15327058 | 13554955 | 11675905 | 7892718 |
| 3.1 | 12759844 | 10309760 | 10097375 | 6324673 |
| 3.2 | 9813965 | 7695683 | 7822262 | 4381483 |
| 3.3 | 7353146 | 6167479 | 5421169 | 2298256 |
| 3.4 | 5097398 | 3848457 | 3154951 | 433421 |
| 3.5 | 2965891 | 1439517 | 1111811 | 53227.2 |
| 3.6 | 594396 | 200530 | 148908 | 17732.9 |
| 3.7 | 64921.1 | 29603.4 | 26339.5 | 10654.1 |
| 3.8 | 24008.5 | 15657.3 | 15684.1 | 7745.03 |
| 3.9 | 15829 | 10040.8 | 8507.57 | 6036.91 |
| 4 | 10673.8 | 7789 | 6772.47 | 5042.04 |
| 5 | 2897.98 | 2767.6 | 2662.29 | 2239.96 |
| 6 | 2124.74 | 1955.02 | 1909.88 | 1720.4 |

3F4AP Gran plot

Vi * 10pH

| Vi (ml) | Run 1 |
| --- | --- |
| 1 | 103960973 |
| 2 | 66475643 |
| 3 | 38584778 |
| 3.5 | 26308422 |
| 4 | 13013670 |
| 4.1 | 10801523 |
| 4.2 | 8495145 |
| 4.3 | 6484908 |
| 4.4 | 4533309 |
| 4.5 | 2734043 |
| 4.6 | 765213 |
| 4.7 | 110024 |
| 4.8 | 39861.3 |
| 4.9 | 24339.4 |
| 5 | 15677.2 |
| 5.5 | 6725.09 |
| 6 | 4675.29 |
| 6.9 | 3327.51 |

| Vi (ml) | Run 2 |
| --- | --- |
| 1 | 93938583 |
| 2 | 58938275 |
| 3 | 33162099 |
| 3.5 | 21845273 |
| 4 | 10577977 |
| 4.1 | 9064662 |
| 4.2 | 7488438 |
| 4.3 | 5605872 |
| 4.4 | 3700088 |
| 4.5 | 2001149 |
| 4.6 | 560655 |
| 4.7 | 69507.2 |
| 4.8 | 32867.8 |
| 4.9 | 19225.3 |
| 5 | 13325.2 |
| 5.5 | 6137.32 |
| 6 | 4116.2 |

| Vi (ml) | Run 3 |
| --- | --- |
| 1 | 89539912 |
| 2 | 59559172 |
| 3 | 35107455 |
| 3.5 | 23433726 |
| 4 | 12988980 |
| 4.1 | 11276406 |
| 4.2 | 9464799 |
| 4.3 | 7895709 |
| 4.4 | 5877083 |
| 4.5 | 3467667 |
| 4.6 | 2045097 |
| 4.7 | 695142 |
| 4.8 | 89680.6 |
| 4.9 | 37547 |
| 5 | 21887.2 |
| 5.1 | 13412.2 |
| 5.5 | 5706.61 |
| 6 | 4220.61 |

| Vi (ml) | Run 4 |
| --- | --- |
| 1 | 69976825 |
| 2 | 45117696 |
| 3 | 25286294 |
| 3.5 | 15972606 |
| 4 | 7290546 |
| 4.1 | 5797495 |
| 4.2 | 4252989 |
| 4.3 | 3014365 |
| 4.4 | 1485706 |
| 4.5 | 381203 |
| 4.6 | 69704.7 |
| 4.7 | 27426.1 |
| 4.8 | 14392.5 |
| 4.9 | 10741.6 |
| 5 | 8879.85 |
| 5.5 | 4750.88 |
| 6 | 3303.22 |

**Figure S4.** Gran plot of 3F4AP titrated with HCl.

4AP Gran plot

Vi * 10pH

| Vi (ml) | Run 1 |
| --- | --- |
| 1 | 5310492141 |
| 2 | 3956456305 |
| 3 | 1682396296 |
| 4 | 485748107 |
| 4.1 | 392598068 |
| 4.2 | 254574176 |
| 4.3 | 152165528 |
| 4.4 | 43584788 |
| 4.5 | 673205 |
| 4.6 | 30232.5 |
| 4.7 | 11496.8 |
| 4.8 | 7612.56 |
| 4.9 | 5568.3 |
| 5 | 5208.21 |
| 6 | 4634 |
| 7 | 1882.34 |

| Vi (ml) | Run 2 | Run 3 | Run 4 |
| --- | --- | --- | --- |
| 1 | 5226233625 | 5237358228 | 5223203745 |
| 2 | 3246014386 | 3007461366 | 3398135327 |
| 3 | 1671577009 | 1064633642 | 1370764014 |
| 4 | 565411567 | 297382137 | 288284421 |
| 4.1 | 462022043 | 257010682 | 245413993 |
| 4.2 | 358760592 | 228695706 | 224520102 |
| 4.3 | 248349396 | 189585599 | 187713949 |
| 4.4 | 186348365 | 129306341 | 140247430 |
| 4.5 | 72635941 | 36534875 | 36656054 |
| 4.6 | 6173143 | 356613 | 360209 |
| 4.7 | 175568 | 27508.5 | 30380.4 |
| 4.8 | 27700.8 | 11934.4 | 15317 |
| 4.9 | 11369.6 | 6425.15 | 8612.62 |
| 5 | 8882.25 | 5452.48 | 7329.97 |
| 5.5 | 5489.03 | 4188.35 | 5358.05 |
| 6 | 2767.59 | 2271.7 | 2111.46 |
| 7 | 1991.31 | 1589.77 | 1199.28 |

**Figure S5.** Gran plot of 4AP titrated with HCl.
